# Brassinosteroid and gibberellin signaling are required for Tomato internode elongation in response to low red: far-red light

**DOI:** 10.1101/2024.02.29.582690

**Authors:** Linge Li, Jesse Wonder, Ticho Helming, Gijs van Asselt, Chrysoula K. Pantazopoulou, Yorrit van de Kaa, Wouter Kohlen, Ronald Pierik, Kaisa Kajala

**Author notes:** **Email address for each author:**.

## Abstract

In this study, we explore the dynamic interplay between the plant hormones gibberellins (GA), brassinosteroids (BR), and Indole-3-Acetic Acid (IAA) in their collective impact on plant shade avoidance elongation under varying light conditions. We focus particularly on low Red: Far-red (R:FR) light conditions achieved by supplementing the background light with FR. Our research delves into how these hormones individually and synergistically influence stem elongation in tomato plants. Through meticulous experimental modulations of GA, IAA, and BR, we demonstrate that GA and BR are sufficient but also necessary for inducing stem elongation under low R:FR light conditions. Intriguingly, while IAA alone shows limited effects, its combination with GA yields significant elongation, suggesting a nuanced hormonal balance. Furthermore, we unveil the complex interplay of these hormones under light with low R:FR, where the suppression of one hormone’s effect can be compensated by the others. This study provides insights into the hormonal mechanisms governing plant adaptation to light, highlighting the intricate and adaptable nature of plant growth responses. Our findings have far-reaching implications for agricultural practices, offering potential strategies for optimizing plant growth and productivity in various lighting environments.

**Highlight:** This study unveils the interplay of brassinosteroids and gibberellins in shade avoidance elongation, revealing how tomatoes acclimate in response to far-red enriched light conditions.

## Introduction

The agricultural sector faces significant challenges with a rapidly increasing global population and a finite amount of arable land. The escalating demand for food, fuel, and fiber necessitates higher yields from field crops, a need not fully met by current production methods (Smith and Whitelam, 1997). To address this, farming practices have shifted towards cultivating plants in dense vegetation to maximize yield from the limited land area. While optimizing land use, dense vegetation introduces its own set of problems. It increases competition among plants for essential resources, notably light. This competition is exacerbated by shade conditions within dense canopies, characterized by an enrichment of far-red (FR) light and a reduction in overall light at increasing planting density (Pierik and De Wit, 2014; Courbier *et al*., 2021; Huber *et al*., 2021; Pantazopoulou *et al*., 2021).

Plants react to these light conditions through Shade Avoidance Syndrome (SAS), a series of developmental responses triggered by the altered red (R) to far-red (FR) light ratio (R:FR) and blue light depletion (Ueoka-Nakanishi *et al*., 2011; Keuskamp *et al*., 2012; Abenavoli *et al*., 2016). Phytochromes and cryptochromes photoreceptors, play a crucial role in detecting these changes and initiating SAS, leading to adaptations such as increased height and altered leaf architecture (Bush *et al*., 2015; Courbier *et al*., 2020; Küpers *et al*., 2023). Like other dicotyledonous plants, tomato plants *(Solanum lycopersicum*) exhibit varied responses to SAS. While stem elongation is a common SAS feature, the underlying cellular mechanisms, particularly in tomato, remains poorly understood (Bush *et al*., 2015; Courbier *et al*., 2020; Spaninks *et al*., 2020).

The stem, a critical organ for nutrient transport and support, undergoes primary and secondary growth phases crucial for plant development (Sanchez *et al*., 2012; Tonn and Greb, 2017). The plant hormone auxin plays a dominant role in stem growth and secondary growth together with strigolactone and jasmonate signaling (Perrot-Rechenmann, 2010; Sehr *et al*., 2010; Agusti *et al*., 2011). SAS is predominantly governed by hormonal regulation, with important roles for auxin, gibberellins (GAs), and brassinosteroids (BRs), modulating plant growth and development in response to light conditions (Bou-Torrent *et al*., 2014; Colebrook *et al*., 2014; Xu *et al*., 2014; Kohnen *et al*., 2016; Iglesias *et al*., 2018). The hormonal changes are triggered by light perception by the photoreceptors. Particularly, low R:FR light conditions are perceived by phytochromes and prompt various developmental changes, including stem and petiole elongation, and reduced branching (Yang and Li, 2017). Research primarily in Arabidopsis, and more recently in legumes like soybean, has shown that low R:FR conditions induce the expression of auxin biosynthesis and transport genes, contributing to stem elongation (Keuskamp *et al*., 2010*b*; Ueoka-Nakanishi *et al*., 2011; Procko *et al*., 2016). Phytochromes are light receptors that suppress PIFs (PHYTOCHROME INTERACTING FACTOR), a family of transcription factors that orchestrate gene expression to regulate processes like chloroplast development, hormone production and cell elongation in ArabidopsiS (Leivar *et al*., 2012; Kohnen *et al*., 2016; Yang *et al*., 2021). PIFs engage with signaling pathways, including auxin and gibberellins, to coordinate plant growth and development in response to environmental signal (Casal, 2013; de Lucas and Prat, 2014; Ma and Li, 2019). Several PIFs can bind to the promoters of *YUCCA* genes in Arabidopsis, which encode enzymes involved in auxin biosynthesis (de Lucas and Prat, 2014; Kohnen *et al*., 2016; Müller-Moulé *et al*., 2016).

Auxin is perceived by TIR1/AFB receptors, and its signaling involves AUXIN RESPONSE FACTORs (ARFs) and Aux/IAA proteins, whose interplay is modulated by light through cryptochrome 1 (CRY1) and phytochrome B interaction by direct inhibition of auxin signaling (Stacey *et al*., 2016; Luo *et al*., 2018; Xu *et al*., 2018). Specifically, ARFs such as ARF7 and ARF19, are crucial for the expression of genes that promote cell elongation in response to shade (Okushima *et al*., 2007). In SAS, GAs also play a critical role, with their levels and distribution being modulated by PIFs under shade conditions, impacting DELLA proteins and thus regulating SAS-related growth processes (Djakovic-Petrovic *et al*., 2007; De Lucas *et al*., 2008; Feng *et al*., 2008; Küpers *et al*., 2023).

Despite the known importance of hormonal regulation in stem growth, the specifics of these mechanisms in the context of SAS in tomato are not fully understood, especially in younger plants (Cagnola *et al*., 2012; Courbier *et al*., 2020). Our study aims to fill this gap by examining stem-specific developmental changes and identifying transcriptome patterns during low R:FR-induced shade avoidance in tomato. We then explored the hormonal interactions of auxin, GA and BR in tomato SAS through a pharmacological approach. Overall, we provide a deeper understanding of the molecular mechanisms governing plant growth and development in low R:FR environment.

## Materials & methods

### Plant materials and growth conditions

We initiated the germination of *Solanum lycopersicum* seeds (cv Moneymaker, sourced from Intratuin B.V) by placing them in sealed plastic containers lined with dampened paper towels soaked in tap water for a duration of one week. Then, uniform-sized seedlings were transplanted into 7 cm square pots containing meticulously sieved Primasta® soil. The seedlings were cultivated under controlled environmental conditions (maintaining a temperature of 20°C, a relative humidity of 70%, and a photoperiod is of 16 h of light,followed by 8 h of darkness). The initial phase involved seven days of growth under standard white light conditions (WL) obtained from Valoya NS1 spectrum, with a photosynthetically active radiation (PAR; 400 -700 nm waveband) intensity of 200 µmol m⁻² s⁻¹. Subsequently, the seedlings were randomly divided into two experimental groups: white light (WL) and far-red supplementation (WL+FR) where WL spectrum was supplemented with additional FR light (peak emission around 730 nm from Valoya FR-only LEDs) to achieve a specific red-light to far-red light ratio (R:FR) of 0.20.

### Tissue harvest for RNA-seq

The first internode, located between the cotyledons and the first true leaf, was selected as the target tissue. Six internodes from each treatment group were collected into liquid nitrogen at specific time intervals of 6, 24, 30, and 48 hours following the commencement of the FR treatment. In total, we amassed a dataset consisting of 32 individual samples, encompassing two light treatments (WL and WL+FR), four distinct time points, each group with three to four biological replicates.

### RNA-seq library preparation

We prepared RNA-seq libraries according to the random-primer primed method published originally in (Townsley *et al*., 2015) with modifications from Kajala et al., 2021. We diverged from the protocol at the final steps, specifically: We cleaned the enriched libraries with 1.1 volume of well-resuspended Ampure beads. One μl of the samples were used for quality check by Bioanalyzer High Sensitivity DNA Analysis kit, and we selected cleaned library of 200-350 bps based on the peaks. Libraries were pooled together for sequencing with the same target concentration. A final clean-up was done with 0.8 volumes of Ampure beads and elution to the final target volume.

### Sequencing and alignment-

Barcoded libraries were sequenced on the High Output: 1 x 75 bp Illumina NextSeq500 at USEQ (Utrecht Sequencing Facility). The initial read quality assessment with FastQC, followed by sequence trimming using TrimGalore (https://github.com/FelixKrueger/TrimGalore/blob/master/Docs/Trim_Galore_User_Guide.md), rRNA removal via SortMeRNA, read mapping and read-group annotation utilizing STAR (Dobin *et al*., 2013), alignment quality control with RSeQC (Wang *et al*., 2012) and Preseq ( https://smithlabresearch.org/software/preseq/), PCR duplicate detection employing Sambamba MarkDup (Tarasov *et al*., 2015), and gene expression and biotype quantification through featureCounts (Liao *et al*., 2014). The sequences were mapped against ITAG4.1 annotation of tomato reference genome SL4.1 (https://solgenomics.net/organism/Solanum_lycopersicum/genome).

### Differential expression analysis

We adapted the DE analysis method from Kajala *et al*., 2021. Libraries with at least 500,000 mapped raw exon counts were selected for analysis. First, we wanted to identify the differentially expressed genes (DEGs) for various comparisons. Raw counts were first converted to count per million (CPM) using with edgeR package (Robinson *et al*., 2009). Genes with CPM > 0.5 in at least one sample for all 4 biological replicates were kept for the analysis. CPM values were quantile normalized with the voom function (Law *et al*., 2014). Principal component analysis (PCA) was conducted to distinguish the separation of samples by timepoint, treatment and cultivar with the ggplot2 package (Wickham, 2009) in R.4.1.3 (R Development Core Team, 2010). DEGs were detected with the limma R package (Ritchie *et al*., 2015). The data underwent a Log2 transformation, and a significance threshold of FDR-adjusted p-value (adj.P.Val) ≤ 0.05 was applied to identify differentially expressed genes. The adjusted p-value list of DEGs was obtained for further analysis. Venn plots were generated by DEGs using imageGP website (Chen *et al*., 2022).

### Gene ontology enrichment analysis

We performed Gene Ontology (GO) enrichment analyses using the GOseq R package (Young *et al*., 2010). GO annotations (ITAG4.1) (Hosmani *et al*., 2019)were obtained from Sol Genomics Network (solgenomics.net). Significantly enriched terms had a p-value < 0.05 and a fold enrichment > 1. Visualization was done using R’s ‘pheatmap’ function.

To expand the annotation for hormones, we looked for Arabidopsis GO annotations (Table 1). We sourced data from TAIR 10, extracted genes responsive to GA, IAA, and BR, and annotated the best tomato matches through a multi-blast on Solgenomics with corresponding TAIR gene names.

**Table 1.**
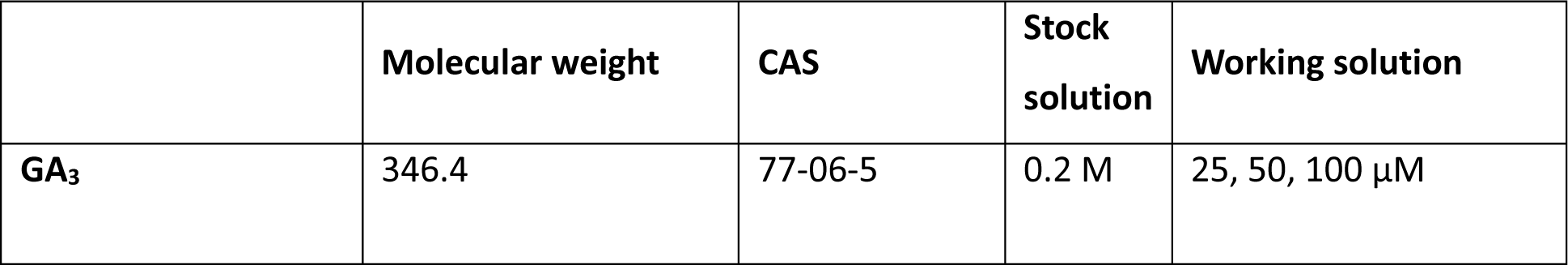
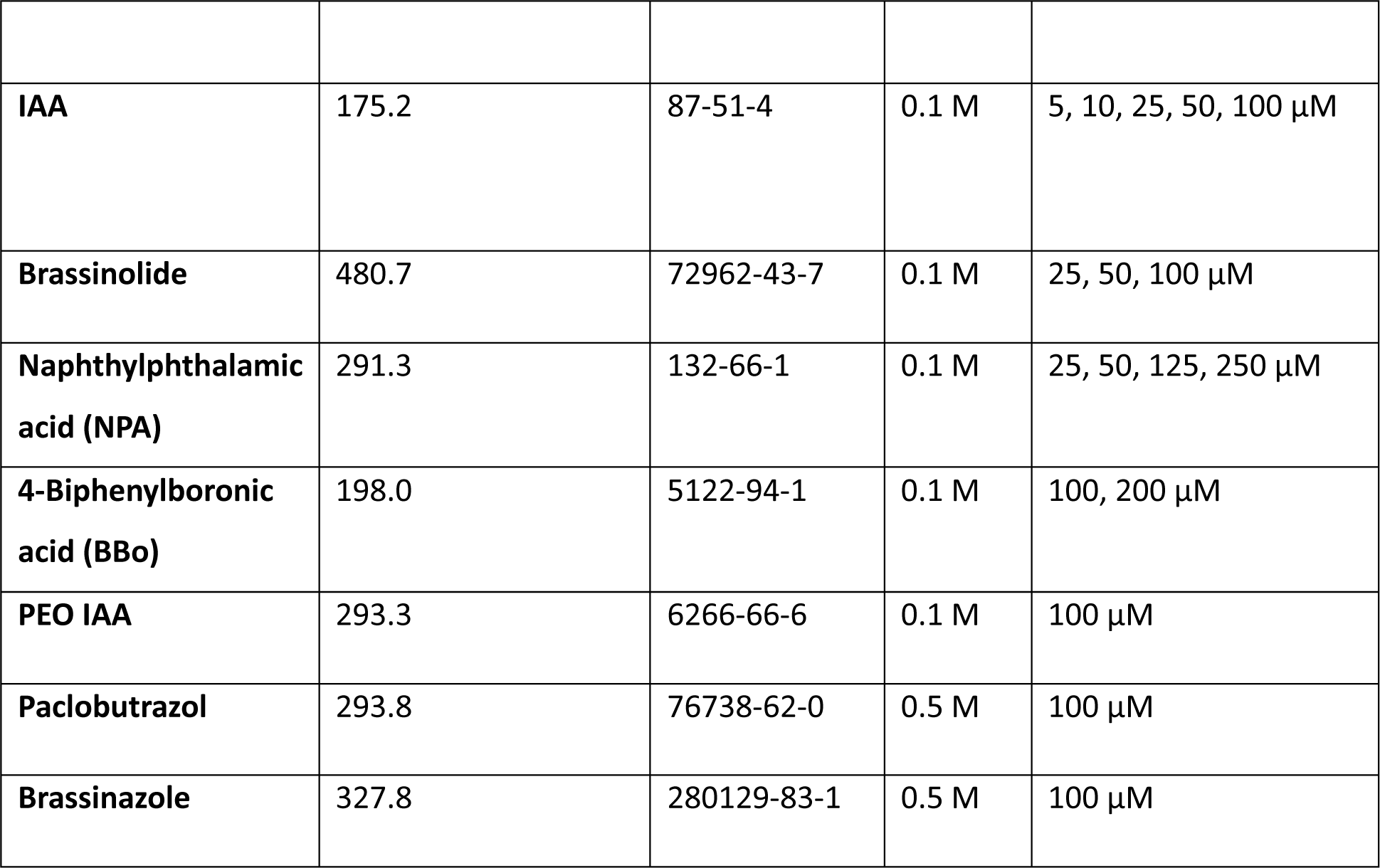
Plant hormones and inhibitors used in this study.

|  | Molecular weight | CAS | Stock solution | Working solution |
| --- | --- | --- | --- | --- |
| GA<sub>3</sub> | 346.4 | 77-06-5 | 0.2 M | 25, 50, 100 µM |
| **IAA** | 175.2 | 87-51-4 | 0.1 M | 5, 10, 25, 50, 100 µM |
| **Brassinolide** | 480.7 | 72962-43-7 | 0.1 M | 25, 50, 100 µM |
| **Naphthylphthalamic acid (NPA)** | 291.3 | 132-66-1 | 0.1 M | 25, 50, 125, 250 µM |
| **4-Biphenylboronic acid (BBo)** | 198.0 | 5122-94-1 | 0.1 M | 100, 200 µM |
| **PEO IAA** | 293.3 | 6266-66-6 | 0.1 M | 100 µM |
| **Paclobutrazol** | 293.8 | 76738-62-0 | 0.5 M | 100 µM |
| **Brassinazole** | 327.8 | 280129-83-1 | 0.5 M | 100 µM |

### Pharmacological treatments

Hormone and hormone inhibitor treatments (Listed in table 1) were initiated once the seedlings reached an appropriate size, approximately 14 days after germination, for inhibitor application, 2 day younger plants were used. For plants exposed to the white light (WL) + far-red (FR) treatment, they were transitioned to a R:FR of 0.2 on the first day of treatment. All measurements were taken on the seventh day post-treatment to evaluate the impact of the hormone and light treatments on the specified plant parameters.

We used multiple chemicals explore the influence of plant hormones and their inhibitors on tomato growth and development under varying light conditions (Table 2) (Kamoutsis *et al*., 1999; Asami *et al*., 2000; Hayashi *et al*., 2008; Mashiguchi *et al*., 2011; Kakei *et al*., 2015). To examine the impact of chemical treatments, 14-day-old tomato plants received working concentrations of the chemical applied through brushing on the first internode 1, using 350 μL volume per treatment. Hormones were administered on days 1, 3, and 5 of the treatment to the entire first internode of each plant. Hormone inhibitors, on the other hand, were applied either directly on the internode one day before the FR treatment or to the soil using a 40 mL of 100 μM working solution two days before the FR treatment. Additionally, a homogeneous spray of the entire plants was tested for 100 μM IAA and 100 μM BBo.

**Table 2.** GO categories used to identify Arabidopsis hormone responsive genes.

| **GO term** | **annotation** |
| --- | --- |
| GO:0009733 | response to auxin |
| GO:0009739 | response to gibberellin |
| GO:0009735 | response to cytokinin |
| GO:1902347 | response to strigolactone |
| GO:0009741 | response to brassinosteroid |
| GO:0009723 | response to ethylene |
| GO:0009751 | response to salicylic acid |
| GO:0009753 | response to jasmonic acid |

### Extraction and Quantification of IAA through Liquid Chromatography-Tandem Mass Spectrometry

To extract IAA from internode tissues, approximately 20 mg of snap-frozen material was used per sample. The tissue was ground manually at -80°C using mortar and pestle. Subsequently, IAA was extracted from these samples overnight at 4C using 1 mL methanol containing [phenyl ^13C6]-IAA (concentration of 0.1 nmol/mL) as an internal standard as previously described (Schiessl *et al*., 2019) (*Schiessl et al., 2019*). After extraction, the samples were filtered through a 0.45 µm Minisart SRP4 filter and analyzed on the same day. The IAA content was quantified using a Waters Xevo TQs tandem quadrupole mass spectrometer, following methods described in previous literature (Gühl *et al*., 2021; Küpers *et al*., 2023).

### RNA extraction, cDNA synthesis & Quantitative RT-PCR

In the qRT-PCR experiment, plants were treated with mock, GA, IAA, GA + IAA, BR, or IAA+GA+BR as described above, and placed under either WL. We sampled four biological replicates, for one biological replicate, a minimum of three internodes were placed in a tube. Plant material was collected 2 hours after pharmacological treatment. All samples were rapidly frozen using liquid nitrogen and stored at - 80°C. Total RNA extraction was performed using the RNeasy kit (Qiagen) according to the manufacturer’s instructions. Reverse transcription was carried out using random hexamer primers with RevertAid H Minus (Thermo Fisher Scientific™) in accordance with the manufacturer’s instructions. We conducted quantitative reverse transcription PCR (qRT-PCR) with three technical replicates for each sample using SYBR Green super mix dye (Thermo Fisher Scientific™) and the CFX Opus 384 and Bio-Rad CFX Maestro platforms. The primers we used are listed in Table S4. Relative transcript abundance was determined through the comparative 2^-ΔΔCt^ method, with *ACTIN2* serving as the reference gene. The data was analyzed using R version 4.3, which included an ANOVA followed by a Tukey test for statistical analysis. Data visualization was performed using GraphPad (Version 9.5.1 (528), January 24, 2023).

## Results

Our research concentrated on the shoot responses of the tomato cultivar Moneymaker (MM) to low R:FR ratio. Following the transplantation of germinated seedlings into soil and a recovery period of one week, we exposed them to light treatments for seven days, either white light with supplemental far-red (WL+FR) or just white light (WL). At 21 days after germination (dag), we collected and analyzed phenotypic data for MM, as illustrated in Figure 1a. Our shoot architecture phenotyping included measurements of traits such as stem length, hypocotyl length, internode lengths, and stem diameters, as detailed in Figures 1b, 1c, and S1. We observed that far-red light (FR) significantly promotes the elongation of stem and each internode (Figure 1c).

**Figure 1:**
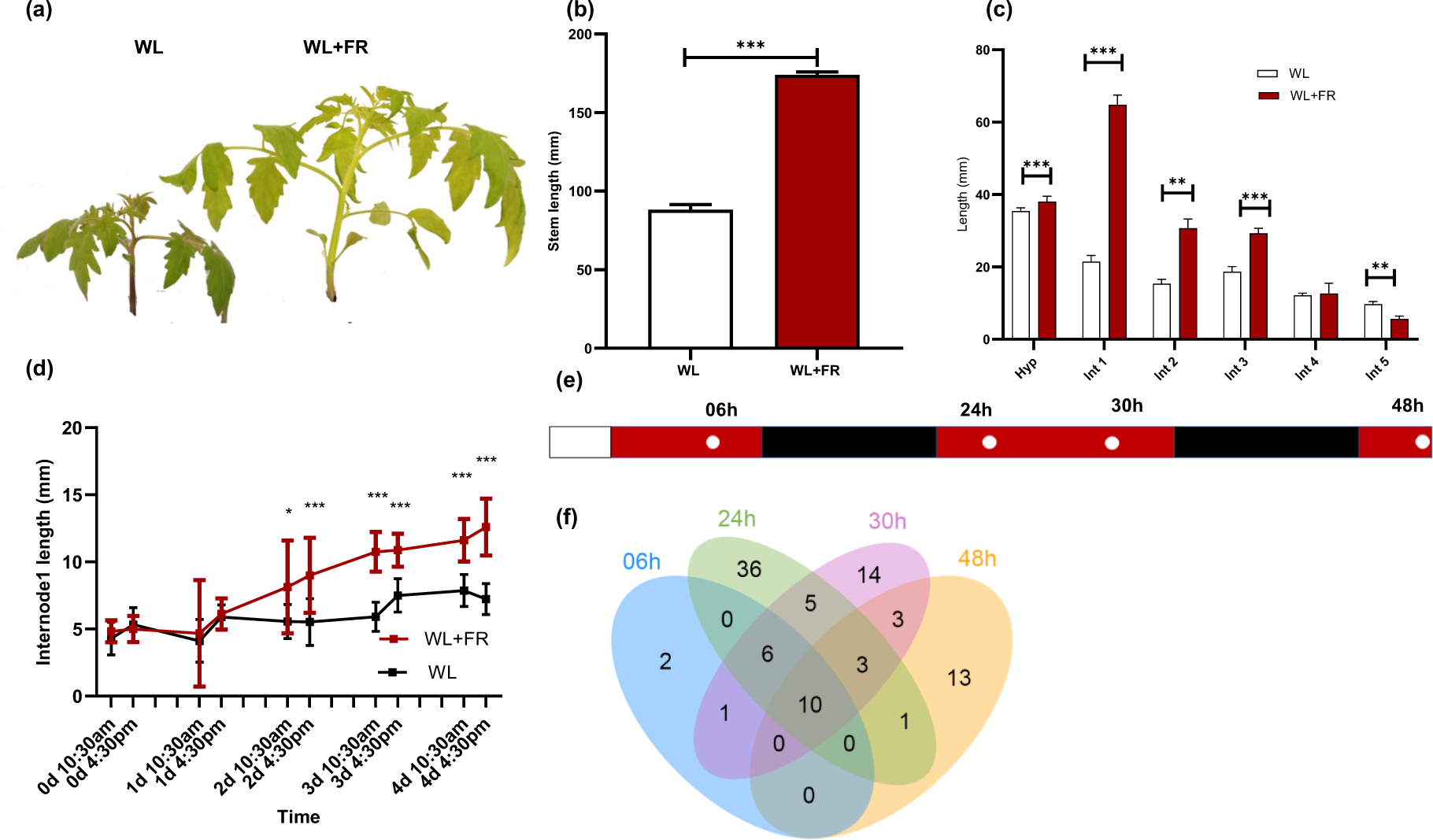
Tomato (MM) stem responses to low R:FR. (a) Comparison of *S. lycopersicum* cultivar Moneymaker 24-day-old young plants in WL (left) vs WL+FR (right). This picture was taken at 10 days of treatment to illustrate the different phenotypes between WL and WL+FR. Hyp=Hypocotyl, Int=Internode. (b) Stem and (c) internode lengths of three-week-old Moneymaker plants after seven-day WL and WL+FR treatments. Biological replicates *n*=18, the experiment was repeated three times. (d) First internode length over 4 days of light treatment. The seedlings were grown in WL for 1 week, and on day 0 at 10:30 am in the morning we started supplemental FR treatment, the lengths were measured at 0, 6, 24, 30, 48, 52, 64, 70, 94 hours. Asterisks indicate significance between WL and WL+FR as follows: * p≤0.05, ** p≤0.01, *** p≤0.001. Experimental design of the time series RNA sequencing. Plants were grown for 7 days in WL before the light treatments. (f) The overlap in FR-responsive DEGs across the timepoints, generated with tool ImageGP (Chen *et al*., 2022).

Distinguishing gene expression in specific tissue is an effective way to reveal tissue functions and tissue-specific regulation. High-resolution gene expression profiling methods to understand SAS have been successfully conducted in Arabidopsis and other models, including tomato (Li *et al*., 2012; Bolger *et al*., 2014; Bush *et al*., 2015; Kohnen *et al*., 2016; Pedmale *et al*., 2016; Gommers *et al*., 2017; Pantazopoulou *et al*., 2017; Molina-Contreras *et al*., 2019; Courbier *et al*., 2020; Küpers *et al*., 2023). We employed high-resolution gene expression profiling to capture the early stages of first internode elongation. Internode lengths were measured with digital caliper every day at 10:30 and 16:30. For one-week FR treatment and pharmacological treatments, internode lengths were measured with digital caliper by the end of treatment at 10:30am. We planned our RNA-sequencing timepoints to coincide with the onset of the elongation phenotype. Samples were collected at 6 hours (16:30 on day 0), 24 hours (10:30 on day 1), 30 hours (16:30 on day 1), and 48 hours (10:30 on day 2), as depicted in Figure 1d. We mapped the reads to the SL4.0 tomato genome (https://solgenomics.net/) using the ITAG4.1 annotation and identified the differentially expressed genes (DEGs) by limma.

First, we examined the overlap of upregulated genes in the internode over time (Figure 1e). Intriguingly, at the 6-hour mark, we observed a relatively low number of DEGs, indicating that the initial response to FR in MM is slower compared to Arabidopsis which responded in 40 mins (Küpers *et al*., 2023). However, as time progressed, an increased number of DEGs emerged at the later timepoints, suggesting a more pronounced response to FR exposure over time.

### IAA and GA-related transcripts show a robust fold change response to FR, while BR-related genes exhibit mild responses

First, we explored the transcriptome changes using Gene Ontology (GO) enrichment analysis, and we found only a handful GO categories enriched, including response to auxin (Figure 2a, Figure S2). We also wanted to explore how genes related to other hormones responded in our RNAseq data, and noted that many hormone GO terms were limited or non-existent in the tomato GO annotation in ITAG4.1. So, we referenced select Arabidopsis GO annotations (Table 2) from The Arabidopsis Information Resource (TAIR 10) (Lamesch *et al*., 2012) to identify hormone-related and -responsive genes, which we used for best homolog search in tomato (https://solgenomics.net/tools/blast/). We also added genes from literature on tomato hormones (Li *et al*., 1994; Knauss *et al*., 2003; Wu *et al*., 2012; Stortenbeker and Bemer, 2019). Then, we visualized the response of these genes to FR light in our transcriptome data (Figures 2b-c, S3). We observed that genes related to auxin and gibberellins (GAs) demonstrated significant fold changes in response to FR, particularly noted in the *SAUR* (*SMALL AUXIN UP-REGULATED RNA*) and *GA OXIDASE* genes. In contrast, genes associated with brassinosteroids (BRs) showed a less pronounced response (Figure 2c). *SAUR* genes are known for their role in growth regulation and respond to a range of hormones, not just auxin (Stortenbeker and Bemer, 2019). GA oxidases, including Gibberellin 2-oxidases (GA2oxs) and gibberellin 20-oxidase (GA20ox) modulate GA levels in plants by converting active GAs into inactive forms, thus influencing plant growth dynamics (Lo *et al*., 2018). These findings indicate interplay between light signals and hormone pathways.

**Figure 2:**
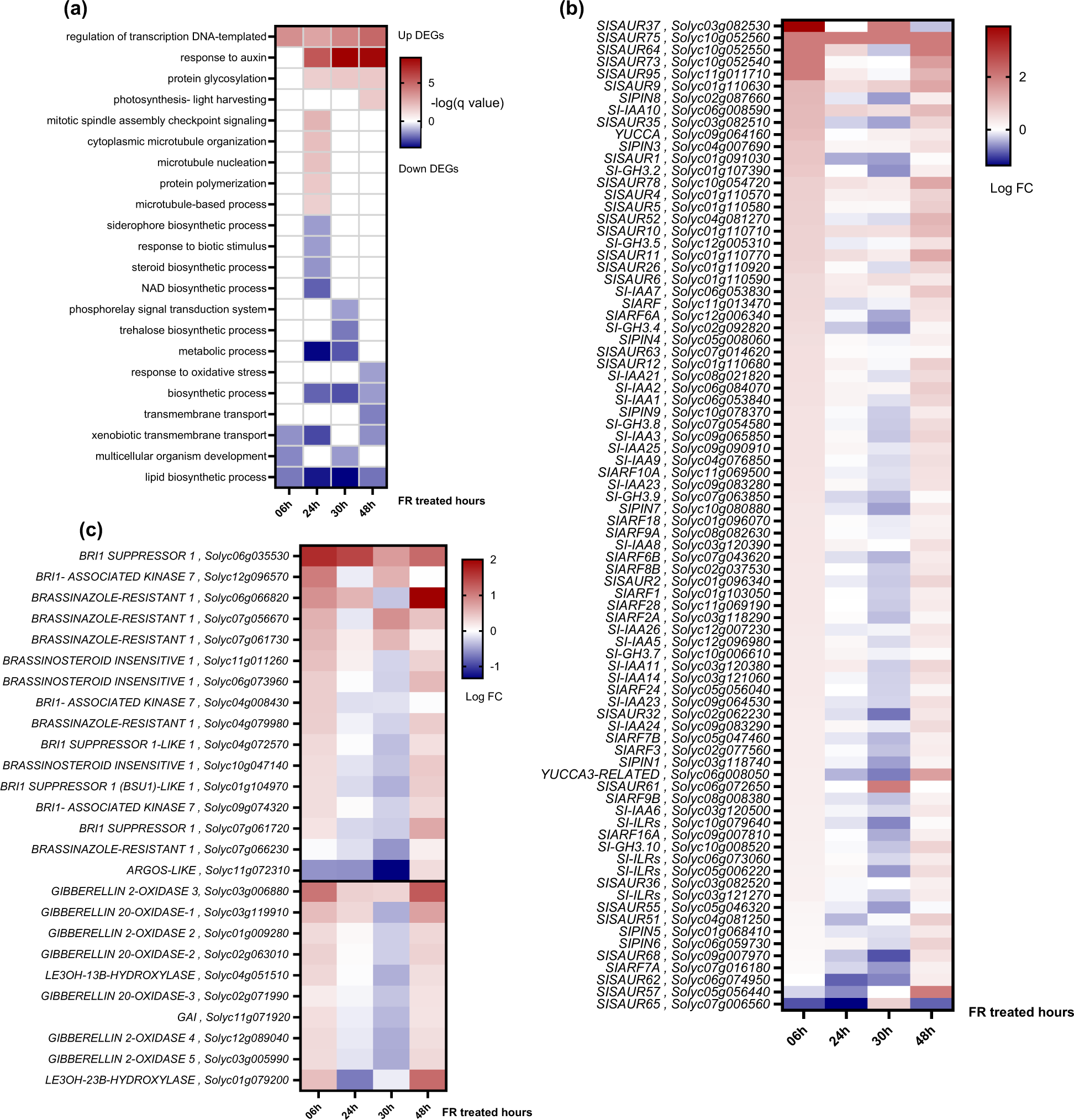
Enrichment of FR light induces transcriptional response to auxin in tomato stems. (**a**) GO enrichment of up- and downregulated DEGs of stems in response to low R:FR. The heatmap displays the negative logarithm (base 10) of the adjusted p-values (q-values) for each enriched GO category, using a color scheme where red indicates upregulated DEGs and blue signifies downregulated DEGs. (**b, c**) FR response in tomato for genes related to auxin, BR, and GA. The heatmaps visualize the log fold changes in gene expression associated with (b) auxin, (c) GA and BR. Red represents upregulation, while blue indicates downregulation in gene expression.

### IAA induced mild stem elongation when applied on first internode or first true leaf regardless of concentration

Next, we wanted to test the role of different hormones for tomato stem elongation. We chose a pharmacological approach using hormone treatments and inhibitors to dissect the hormonal dynamics in SAS. As none of these treatments were previously published for tomato, we first optimized the chemical application methods, timings and frequencies of treatment, and chemical concentrations.

We focused first on the Indole-3-Acetic Acid (IAA)-responsive pathway as suggested by our RNAseq findings (Figure 2a). IAA’s involvement in SAS has been noted in various species, particularly through FR-induced IAA production in leaves, affecting petiole and hypocotyl growth (Procko *et al*., 2014; Kohnen *et al*., 2016; Michaud *et al*., 2017; Pantazopoulou *et al*., 2017; Yang and Li, 2017; Küpers *et al*., 2023). In Arabidopsis SAS research, IAA administration is typically conducted by applying it to different leaf areas or incorporating it into the growth medium (Keuskamp *et al*., 2010*a*; Pantazopoulou *et al*., 2017). However, given the distinct stem-centric growth pattern of tomato as opposed to Arabidopsis’s rosette form, we sought to examine alternative IAA application methods. We experimented with a whole-plant spray and a targeted brushing technique on both the first internode and leaf (Figure 3a,b), aiming to identify the most effective method for probing IAA’s influence on tomato growth. Whole-plant spray did not yield notable responses in the first internode at concentrations of 25 μM and 100 μM (Figure S4), while targeted brushing of 100 μM IAA on the first internode or first leaf led to a slight increase in the length of the internode (Figure 3c).

**Figure 3:**
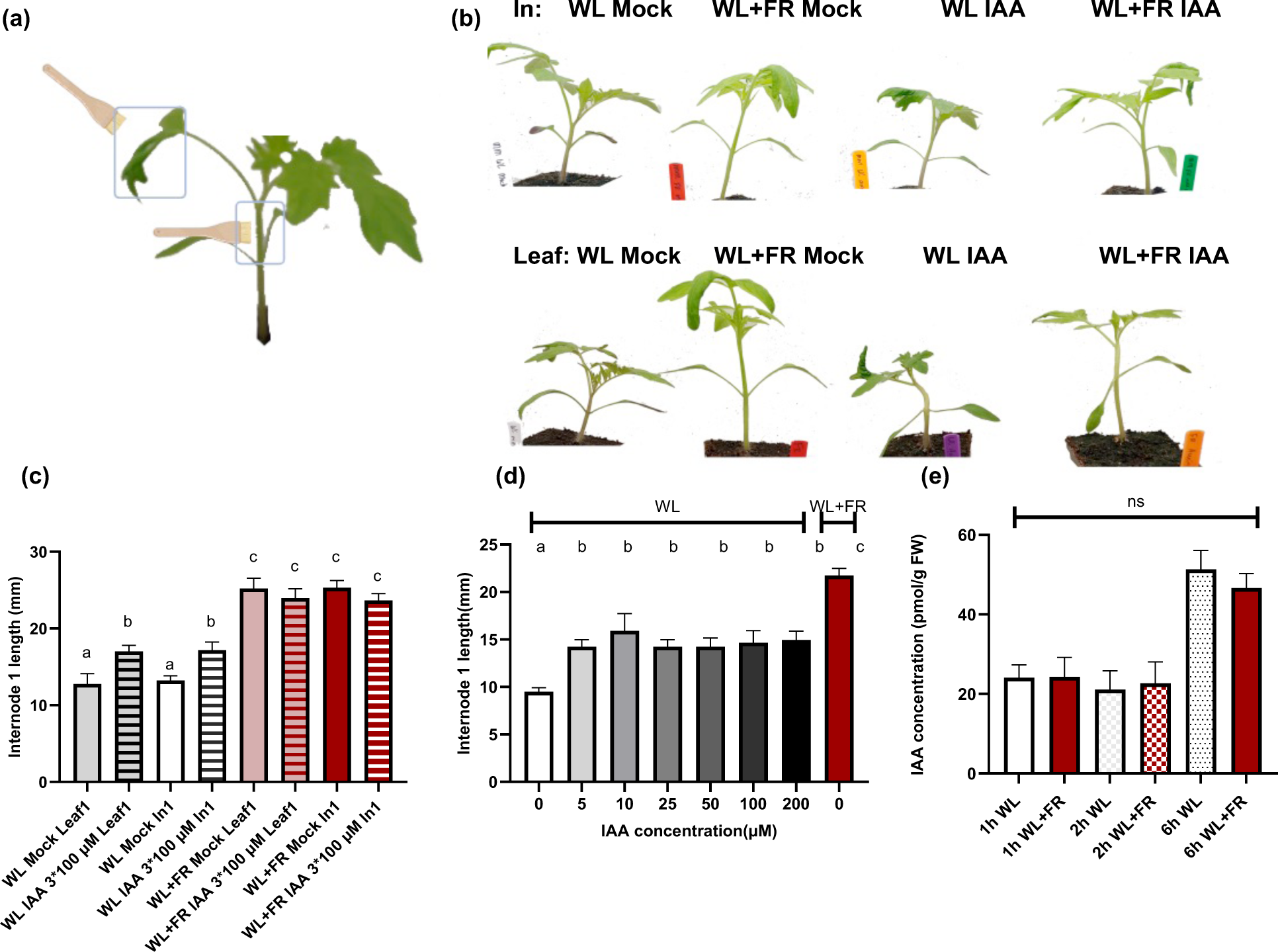
IAA is not sufficient to induce the low R:FR phenotype. (a-d) We treated 14-day-old tomatoes with different concentration IAA by brushing onto internode 1 or leaf 1. IAA or mock were applied 3 times over one week treatment period. There are 12 biological replicates, and the experiment was repeated twice. (a) Illustration of how internode and leaf brushing was carried out. (b) Representative photos of the plants after the treatments. (c) Internode 1 length in different IAA and light treatment combinations. (d) Internode 1 length in response to different IAA concentrations applied to the internode 1. (e) We measured the IAA concentration in internode 1 in response to 1-hour, 2-hour, and 6-hour WL and WL+FR treatments. Data are presented as mean ± SEM, and different letters indicate significant differences between treatments based on ANOVA analysis with Tukey’s post hoc test (P<0.05).

Our aim was to test if we could replicate the effects of FR treatment using localized IAA applications. Applying 100 μM IAA to either the first leaf or first internode, with three brush applications over a week in WL, we observed that internode 1 elongated to similar extent by both applications, but the elongation did not reach the level induced by FR (Figure 3c). When we tested the effect of IAA in WL+FR, we observed no significant IAA-induced alterations regardless of the application site (Figure S4, S5).

Subsequently, we wanted to determine whether the concentration of IAA applied might be either insufficient or excessive to achieve the full extent of elongation in internode 1, akin to what is observed under FR conditions. To this end, we experimented with a spectrum of IAA concentrations, applying them to Internode 1 via brushing, with three applications spread over the duration of a week, mirroring the length of the FR treatment. Contrary to our expectations, the extent of elongation in internode 1 showed no variation in response to different IAA concentrations (Figure 3d). Notably, at both 5 μM and 200 μM concentrations of IAA, the degree of internode 1 elongation was statistically identical. This finding implies that the elongation response of internode 1 to IAA is not dependent on the concentration of IAA used, indicating that even the lowest concentration of IAA we applied was sufficient to trigger the maximum response achievable solely through IAA (Figure 3d, S4, S5). Overall, we concluded that IAA alone was not sufficient to drive internode elongation to the same extent as WL+FR treatment does.

### Auxin levels in the internode were unaffected by FR enrichment

Given the observed induction of auxin response in the transcriptome and the influence of externally applied IAA, we were intrigued to explore if internal IAA levels in the internode change in response to FR treatment. To this end, we utilized liquid chromatography-tandem mass spectrometry for quantifying the IAA concentration within the first internode following FR treatments of 1-hour, 2-hour, and 6-hour durations (Figure 3e). We did not observe any increase in IAA concentration in the WL+FR treatment compared to WL in internode 1. This suggests either a lack of regulation of IAA concentration by FR or that the FR-induced changes to IAA accumulation are mostly about spatial distribution within the internode that cannot be detected in our whole internode sampling.

### Auxin inhibitors do not affect FR-responsive stem elongation

We wanted to test if auxin was necessary for the FR-induced stem elongation, and we chose three different chemicals that affect auxin. First, we conducted an experiment using N-1-naphthylphthalamic acid (NPA), a polar auxin transport inhibitor, to understand the role of leaf-derived auxin in light signal response and plant growth (Zhao *et al*., 2001; Teale and Palme, 2018; Abas *et al*., 2020). We applied NPA at three different concentrations (25 μM, 50 μM, 125 μM) to the first internode of the tomato plants, starting a day before the seven-day light treatments. The application of NPA markedly affected plant morphology, with the higher concentrations having stronger effect on the ability of the plant to stay upright (Figure 4a). We observed that all concentrations of NPA induced stem elongation in WL (Figure 4b) and interestingly, inhibited the stem diameter thickening in both light treatments (Figure 4c), which could explain the fragility observed at higher concentrations.

**Figure 4:**
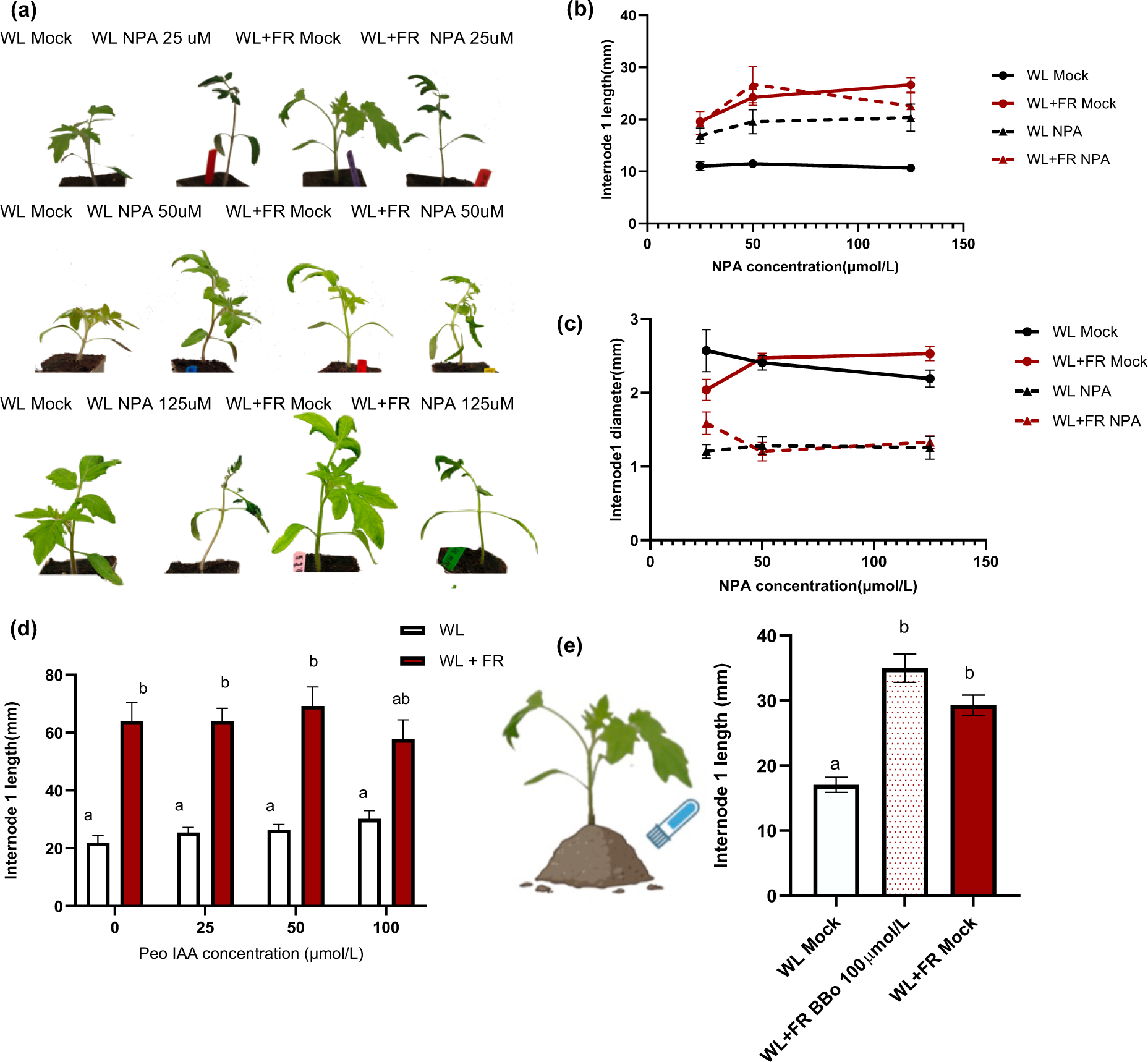
Auxin inhibitors do not suppress low R:FR-induced stem elongation in tomato. (a-c) We treated the first internode of 13-day-old tomatoes with 25, 50, 125 μM NPA (or mock) in WL conditions one day prior to the start of the one-week light treatment. (a) Representative photos of the plants after the NPA and light treatments. (b) Internode 1 elongation and (c) Internode 1 diameter in response to different NPA concentrations and light treatments. (d) First internode length in WL+FR is not affected by different PEO-IAA concentrations. We treated the first internode of 13-day-old tomatoes with different concentration PEO-IAA (or mock) in WL conditions one day prior to the start of the one-week light treatment. (e) First internode length in WL+FR is not affected by 100 μM BBo. We drained the soil 4 days before the light treatment, and applied BBo directly to the soil 2 days before FR treatment in FR conditions. For panels (b-e), data are presented as mean ± SEM, and different letters indicate significant differences between treatments based on ANOVA analysis with Tukey’s post hoc test (P<0.05). There are 12 biological replicates, and the experiment was repeated twice.

Subsequently, we examined the effects of PEO-IAA (2-(1H-Indol-3-yl)-4-oxo-4-phenyl-butyric acid), an auxin antagonist that prevents auxin from binding to its receptor (Hayashi et al., 2008) and 4-biphenylboronic acid (BBo), an IAA biosynthesis inhibitor (Kakei *et al*., 2015). PEO-IAA interacts with TIR1/AFB (TRANSPORT INHIBITOR RESPONSE 1/AUXIN SIGNALING F-BOX) proteins similarly to -alkyl-IAA, and leads to a suppression of auxin-responsive gene expression, cell division, and elongation pathways (Šenkyřík *et al*., 2023). BBo is a chemical that targets YUCCA, the last step in IAA biosynthesis pathway (Kakei *et al*., 2015). We wanted to test if PEO-IAA or BBo would inhibit FR-induced stem elongation, but observed no effect from either compound (Figure 4d,e, S6). This suggests that IAA is not necessary for FR-induced stem elongation in tomato, or that these compounds are not effective in tomato, for example due to differences in IAA pathways or penetration into the relevant tissues.

### GA and BR signaling are necessary for the elongation response induced by FR

As endogenous IAA seemed to have only a limited role in the tomato stem FR-response, we continued our exploration to gibberellins (GAs). Our RNAseq data indicated changes in GA-related gene expression in FR-enriched stems, aligning with known roles of GA in FR-responsive elongation in Arabidopsis (Djakovic-Petrovic *et al*., 2007; Kohnen *et al*., 2016). We wanted to assess the impact of GA and its combined influence with auxin on tomato growth. Bioactive Gibberellic acid 3 (GA_3_) and paclobutrazol (PBZ) are commonly used to study the effects of GAs and their inhibition, respectively, shedding light on GA-mediated growth under shade conditions (Gallardo *et al*., 2002; Desta and Amare, 2021).

Testing various exogenous GA_3_ concentrations revealed a dose-dependent increase in the first internode length in WL (Figure 5a, S7). Under white light (WL) conditions, tomato plants showed progressively increased stem and first internode lengths with higher GA_3_ concentrations, peaking at 100 μM GA_3_ where internode elongation matched that induced by FR light.

**Figure 5:**
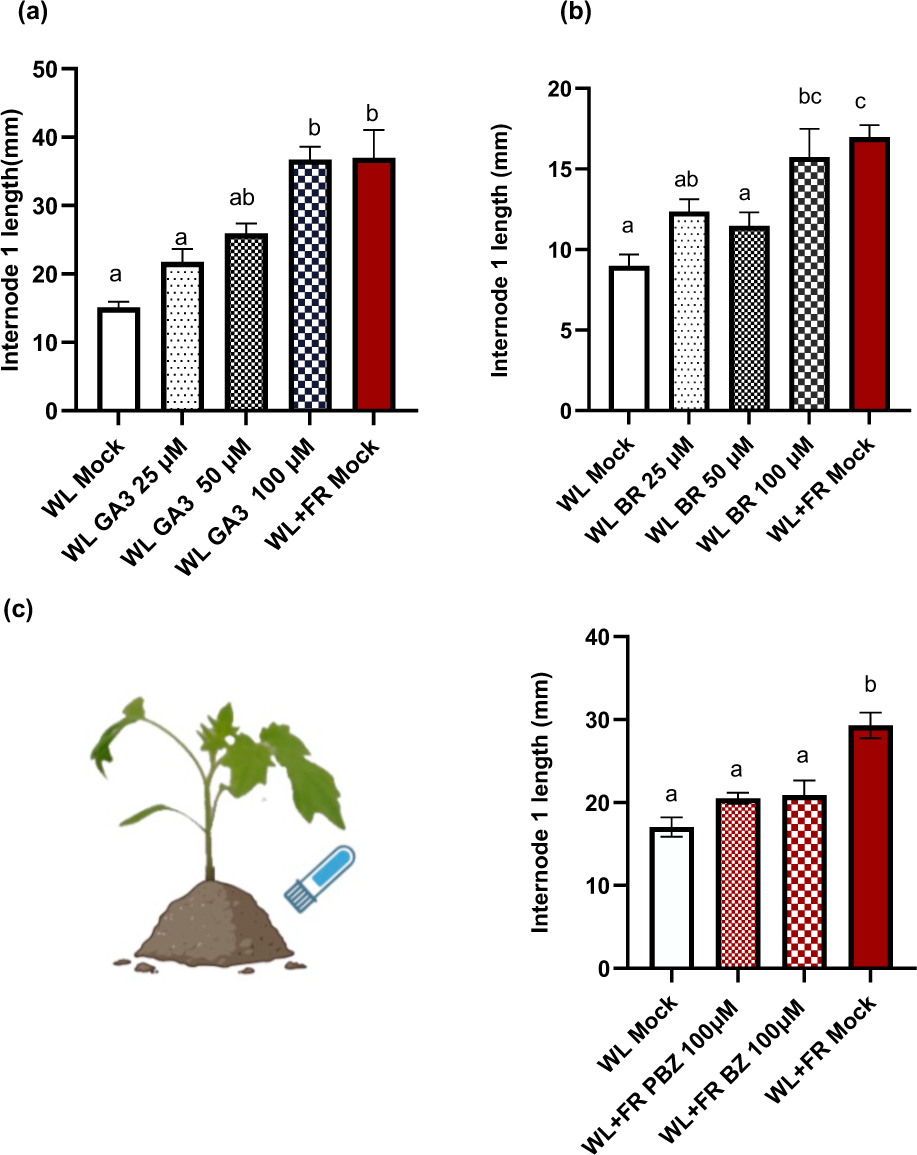
The role of GA and BR in inducing stem elongation under low R:FR conditions. We applied different concentrations of GA_3_ and BR to the first internode of 14-day-old tomato plants. The application, carried out in both white light (WL) spanned one-week light treatment period with brush applications made thrice during this time. (a) Length of the first internode in response to different GA_3_ concentrations. (b) Length of the first internode for the response to different BR concentrations. (c) Length of the first Internode in response to a 100 μM PBZ and BZ in WL+FR. For the inhibitor treatments, the soil was drained four days prior to the FR treatment, and the inhibitors were applied directly to the soil two days before the FR treatment commenced. All data are presented as mean ± SEM, and significant differences between treatments were determined through ANOVA analysis with Tukey’s post hoc test (P<0.05). Each experiment had 12 biological replicates and the entire experiment was repeated twice.

Given that GA_3_ application could mimic FR-induced elongation, we explored whether GA was necessary for this response by using paclobutrazol (PBZ), a gibberellin synthesis inhibitor. We observed that blocking GA synthesis under supplemental FR light resulted in internodes similar to those grown under WL (Figure 5b,S7 a-d), confirming that GA is indeed necessary for FR-responsive stem elongation.

We also examined the role of brassinosteroids (BRs) in this process. Despite only mild effects observed in RNAseq data for BR-related genes, BRs are integral to many growth processes, including cell elongation, which is crucial for hypocotyl elongation (Keuskamp *et al*., 2011; Procko *et al*., 2016). First, we applied Brassinolide, a BR, and observed internode elongation comparable to that seen with FR treatment for 100 μM (Figure 5c, S7e-h). This indicate that also BRs are sufficient to mimic the effect of FR for the internode.

Then, to further understand the role of BRs, we used Brassinazole (BZ), an inhibitor of BR biosynthesis (Asami *et al*., 2000) to test if BRs were necessary for the FR-induced internode elongation. We observed that BZ treatment completely inhibited FR-responsive stem elongation (Figure 5b,S7i-l). These findings collectively show that GAs and BRs are necessary for the FR light-induced internode elongation in tomato, and both can induce internode elongation comparably to FR.

### The combined effects of GA, IAA, and BR in the FR-responsive internode elongation

Next, we wanted to test how these three hormones interacted. First, we tested the synergistic effects of GA, IAA, and BR, starting with the combined application of GA and IAA. While IAA alone had shown only mild effects on elongation, we hypothesized that combining it with GA might enhance this response. We treated plants with both GA_3_ and IAA at 50 or 100 μM, and observed elongation in the first internode length in WL (Figure 6a, S8). Notably, plants treated with both 100 μM GA_3_ and IAA exhibited longer internodes than the WL+FR treated plants. This effect was also seen in WL+FR light, suggesting an additive effect of these hormones and FR supplementation on stem elongation. When we combined GA with BR (Figure 6b, S8), we observed again internode 1 elongation that was stronger than in WL+FR treatment. Finally, we tested the effects of all GA, IAA, and BR combined (Figure 6c, S8). Intriguingly, this trio of hormones induced internode elongation similar to that observed under supplemental FR treatment regardless of the concentration (Figure 6c). This finding contrasts with the results from GA+IAA and GA+BR treatments, where the induced internode elongation exceeded that of FR. This suggests that while GA, IAA and BR might have additive effects on stem elongation, the combination possibly moderates this response.

**Figure 6.**
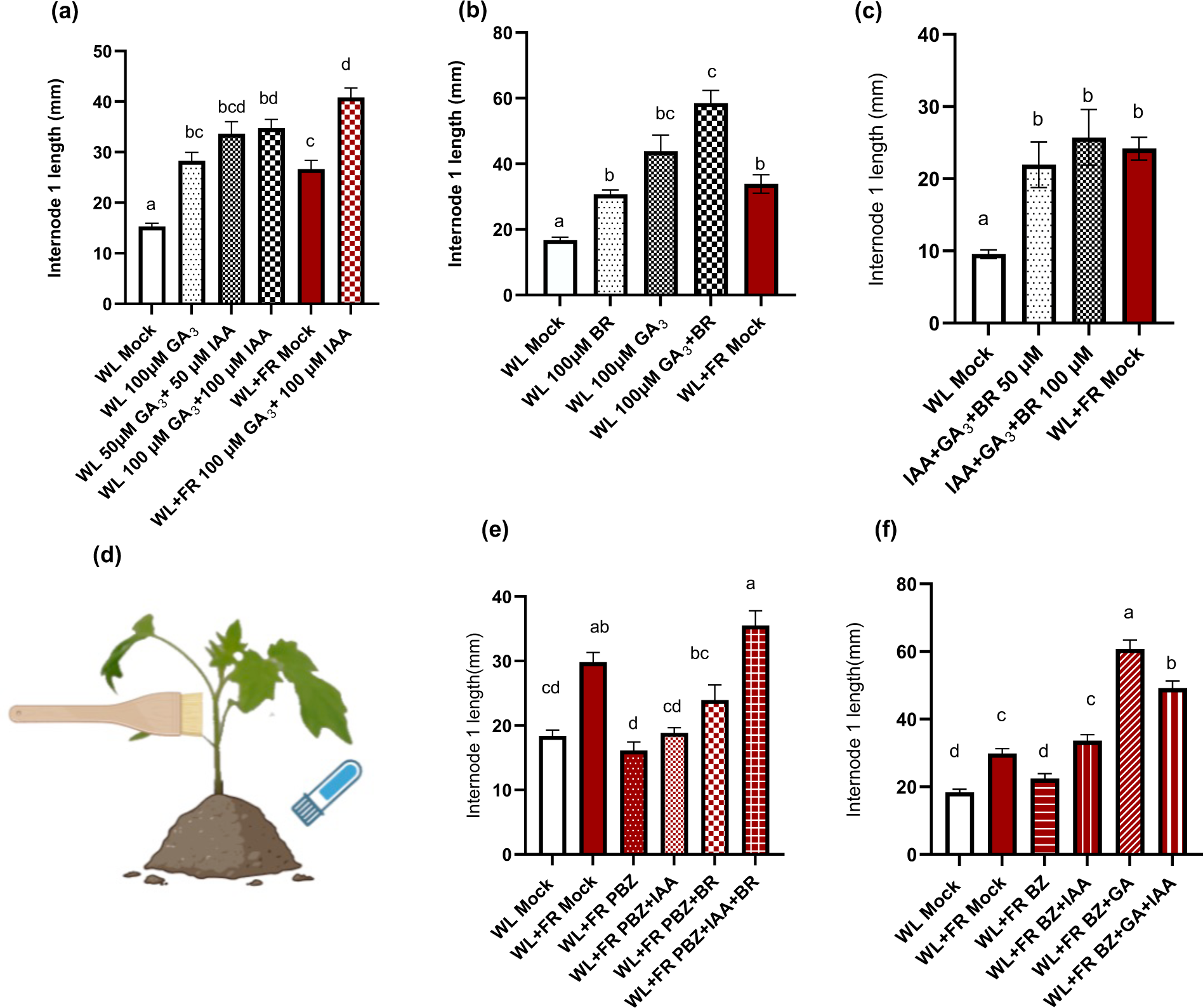
Cross-talk of IAA, BR, and GA in FR-induced internode elongation. (a-c,e-f) Internode 1 lengths of 14-day-old plants in response to hormone treatments applied thrice by brushing on the internode over the one-week light treatment period. (a) Response to 50 μM and 100 μM GA_3_ and IAA combined in WL or WL+FR. (b) Response to 100 μM BR, GA_3_, or BR+GA_3_ in WL. (c) Response to 50 or 100 μM GA+IAA+BR. (d) The inhibitors were applied by soil penetration two days before the one-week light treatment period started, during which we applied hormones (or mock) to the internode by brushing thrice. (e) Response to 100 μM PBZ followed by 100 μM IAA and 100 μM BR in WL+FR. (f) Response to 100 μM BZ followed by 100 μM IAA and 100 μM GA_3_ in WL+FR. Data presented as mean ± SEM, with different letters indicating significant differences between treatments based on ANOVA analysis with Tukey’s post hoc test (P<0.05). Six biological replicates. Picture created by Biorender.

To further study the crosstalk of these hormones, we conducted rescue experiments. We applied either PBZ or BZ in WL+BR to suppress GA or BR biosynthesis and inhibit the FR-responsive internode elongation, and tested if the other two hormones could rescue the phenotype (Figure 6d-e, S7i-l). BR counteracted the effect of GA synthesis inhibition and maintained internode elongation comparable to WL+FR already on its own, and addition of IAA led to a synergistic effect (Figure 6d). Reciprocally, GA application overcame the inhibition of BR biosynthesis and led to a doubling of internode length compared to that induced by WL+FR (Figure 6e,S9). Unlike GA inhibition where IAA addition had no effect, we observed that IAA could rescue the elongation inhibition due to BZ. However, combining BZ with both IAA and BR resulted in an elongation level that was intermediate between the effects of BR or IAA alone, yet still higher than that induced by FR light.

Overall, we show that while GA and BR are necessary for FR-induced internode elongation, their effects seem to be additive, and either hormone can compensate for the absence of the other, also when combined with IAA. However, the role of IAA in this hormonal crosstalk remains complex, potentially influenced by factors like internal concentrations or spatial distribution.

### IAA and GA induced the most similar gene expression changes compared to FR

The ability of IAA, GA and BR to induce internode elongation in tomato does not necessarily imply that they are sufficient to trigger specifically the FR-responsive internode elongation. To gain more insight into the molecular events downstream of the supplementary FR treatment and how the different hormone treatments mimic it, we carried out a quantitative reverse-transcription PCR (qRT-PCR).

We selected genes that were upregulated by at least threefold in response to FR supplementation, as determined by differential expression (DE) analysis (Figure 1), and profiled their expression in the first internode in the following hormone treatments in WL: GA, IAA, BR, GA+IAA, BR+GA+IAA, each at 100 μM, as well as mock treatment in WL and WL+FR (Figure 7a-v) after two hours FR treatment based on a time series qPCR result (Figure S10). None of the hormone treatments stood out as reproducing the pattern of FR-responsive gene expression, so we calculated the Pearson’s correlation coefficients between the samples (Figure 7w). This revealed that the IAA-induced gene was the closest match with FR-induced gene expression. We also noticed that BR treatment elicited a distinct and robust expression profile, diverging significantly from the patterns induced by other hormones and far-red (FR) light. Moreover, the impact of BR was so pronounced that it overshadowed the influences of GA and IAA, as evidenced by the high correlation between the gene expression patterns in BR treatments and those in the combined BR+GA+IAA treatments. Conversely, GA on its own did not resemble the other treatments but was overshadowed by the IAA treatment.

**Figure 7:**
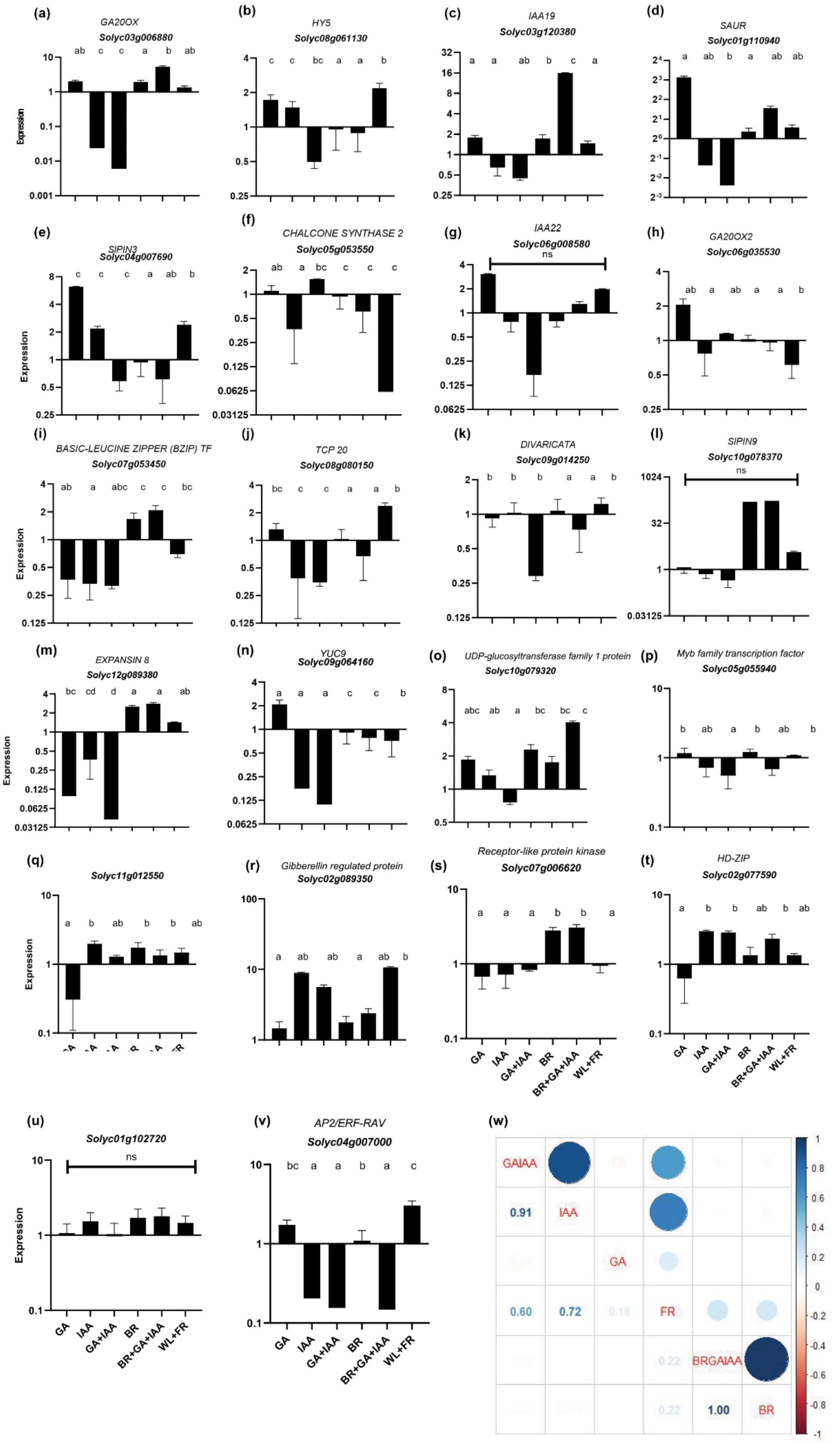
Analysis of FR-responsive gene expression in response to hormone treatments. This figure presents a detailed examination of gene expression in response to hormone treatments and far-red (FR) light, focusing on select genes that were significantly upregulated (fold change >3) in RNA-seq data. We conducted quantitative PCR (qPCR) analysis on samples treated with 100 μM concentrations of GA, IAA, BR, combinations of GA+IAA, BR+GA+IAA, or exposed to FR light. The expression levels were measured in internode after a 2-hour treatment duration. To ensure the robustness and reliability of our findings, the experiment was structured to include four biological replicates, each comprising 8-12 plants. The results are depicted as mean ± SEM. Distinct letters denote significant differences determined from ΔΔCt values by ANOVA followed by Tukey’s post hoc test (P<0.05). Additionally, three technical replicates were performed for each biological replicate to validate the accuracy of the qPCR measurements. An integral part of our analysis was the comparison of hormone induced gene expression changes against FR. This comparison was quantified using Pearson’s correlation matrix (w). Significant differences at p<0.05 among relatedness treatments after post hoc multiple comparison are shown separately for each treatment on the left and representation of the fitness with dot colored in blue for positive correlation, red for negative correlation.

We considered that the target gene selection could be skewing our results towards IAA correlating well with FR. We removed genes known to respond to IAA, GA, or BR from the dataset (Figure S11), the correlations among GA, IAA, and BR with FR treatments became more uniformly distributed, but IAA still remained the top match for FR.

All in all, it appears that while BR and GA are necessary for the FR-induced internode elongation in tomato and can drive similar elongation response, these hormones do not mimic FR-driven gene expression patterns. Intriguingly, our data indicated that IAA was not necessary for FR response or sufficient to drive the full internode elongation, this can also be intrinsically biased by the choice of genes.

## Discussion

Crops frequently grow in environments where resources are under high competition. Especially light becomes a limiting factor in densely planted areas. The Shade Avoidance Syndrome (SAS) is a typical adaptive response observed in many commercial crops, including raspberries, apples, and maize that suffer from light shortage-related yield loss (Maughan *et al*., 2017). Tomato is also one of these plants. In a previous study by Bush et al. (2015), the authors reported shade effects on tomato stem elongation and the shoot apical meristem transcriptome, yet the signaling pathways in tomato underlying this response remained untested. To bridge this knowledge gap we carried out tomato low R:FR treatment to characterize the internode responses and the associated transcriptome changes. Differentially expressed genes (DEGs) of RNA sequencing revealed a lower-than-anticipated number of DEGs (Figure 1e) in comparison to Arabidopsis SAS datasets (Kohnen *et al*., 2016; Küpers *et al*., 2023). The transcriptome changes (Figure 2) indicated a role for auxin signaling, so we followed by with a study of the hormonal dynamics of the first internode elongation in response to low R:FR light in tomato. As the role of auxins (IAA) did not appear to be as comprehensive as in Arabidopsis, we also investigated the roles of gibberellins (GA), and brassinosteroids (BR).

We examined the role of IAA in internode elongation (Figure 3,4, S4, S5). The role of IAA in Arabidopsis shade avoidance is characterized in detail for both hypocotyl elongation of a seedling and petiole hyponasty in a mature leaf (Kohnen *et al*., 2016; Yang and Li, 2017; Du *et al*., 2018). In both cases, IAA is produced distally, in cotyledons and leaves, in response to low R:FR sensed by phytochrome B. The signal is conveyed through PIFs to induce *YUCCA* expression and IAA biosynthesis (Michaud *et al*., 2017; Ma and Li, 2019). IAA is then the long-distance signal to hypootyl or petiole. Our results were inconclusive about the role that IAA plays in tomato SAS. We observed that IAA treatment led to increased elongation in first internode (Figures 3+S3), but it was not comparable to FR-induced elongation. The auxin transport inhibitor NPA, the biosynthesis inhibitor BBo and the auxin antagonist PEO-IAA did not inhibit the FR-responsive internode elongation (Figure 4), as would’ve been expected if IAA was necessary for the FR response. Additionally, we did not observe changes to IAA concentration in the internode (Figure 3e). However, the IAA effects might come from specific localization of the hormone within the internode, which we would not be able to detect by bulk measurements or accurately affect by the hormone and inhibitor treatments. The IAA localization has been shown to be a key regulator of the unequal elongation of the two sides of the petiole during FR-induced petiole hyponasty (Michaud *et al*., 2017; Küpers *et al*., 2023)). After all, the RNAseq data (Figure 2) and the qRT-PCR experiments (Figure 7w) strongly point at strong IAA effect on the gene expression during the low R:FR response in the internode.

Given that auxin alone does not fully replicate the internode elongation observed in tomato shade avoidance, we investigated the potential interplay between IAA and other hormones like GA and BR. Our experiments with GA and BR treatments (Figure 5, 6, S7, S8) demonstrated their strong impact on internode elongation, which was additive and more pronounced than that of IAA or even WL+FR. This raises the possibility that GA and BR might activate distinct molecular pathways or interact with distinct key regulators to elongation (Figures 5). The use of inhibitors such as paclobutrazol (PBZ) and brassinazole (BZ) further highlighted the necessity of each GA and BR in FR-induced elongation (Figure 5). We also observed a compensatory relationship between these hormones in regulating tomato internode elongation under low R:FR conditions, as they could rescue the inhibition of each other (Figure 6). In the rescue experiments (Figure 6), we also observed that IAA restores internode elongation back to FR-like level in plants treated with BZ in WL+FR, indicating that IAA can replace the role of BR in this process. Given that IAA alone cannot induce FR-like elongation in WL, it could indicate that WL+FR treatment triggers a signaling pathway where IAA is downstream of BR, or that IAA requires another FR-triggered parallel pathway to induce the full FR-like internode elongation.

In summary, we show that the different hormone and inhibitor treatments, and their combinations, strongly influenced plant growth, with certain combinations capable of either mitigating or amplifying the effects of shade avoidance responses induced by far-red light. This illustrates the complex interplay of hormones in regulating internode elongation in tomato shade avoidance. These insights are obtained using tomato as a study system but could pave the way for broader applications in agricultural science, particularly in understanding hormonal regulation and its application to specific plant traits.

## Abbreviations

SAS: Shade avoidance syndrome
GA: Gibberellins
BR: Brassinosteroids
IAA: Indole-3-Acetic Acid
R: red light
FR: Far-red light
WL: White light
DEG: Differentially expressed genes
GO: Gene Ontology
NPA: Naphthylphthalamic acid
BBo: 4-biphenylboronic acid
PEO-IAA: 2-(1H-Indol-3-yl)-4-oxo-4-phenyl-butyric acid
PBZ: Paclobutrazol
BZ: Brassinazole

## Supplementary files

Table S1: Raw exon counts of RNA-seq analysis samples.

Table S2: Summary of differentially expressed genes (DEGs).

Table S3: Gene ontology (GO) enrichment analysis for molecular function and cell components.

Table S4: qRT-PCR primer sequences.

Figure S1: Internode 1 diameter in response to low R:FR.

Figure S2: Enrichment analysis of far-red light-responsive differentially expressed genes (DEGs) in gene ontology (GO) categories from cellular components and molecular functions.

Figure S3: FR response in tomato internode for genes related to ethylene, cytokinin, strigolactones, ABA, SA and JA.

Figure S4: Evaluating tomato stem growth under exogenous whole plant 25, 100 μM IAA spray application.

Figure S5: Analysis of tomato stem growth following IAA treatments.

Figure S6: Evaluating tomato stem growth under varied NPA and PEO-IAA concentrations.

Figure S7: Analysis of tomato stem growth under various GA, BR, and inhibitor (PBZ, BBO, BZ) concentrations.

Figure S8: Analysis tomato stem response to IAA, GA, BR, and their combinations.

Figure S9: Tomato stem growth under hormonal and inhibitor treatments in WL or WL+FR.

Figure S10: Gene expression in response to low R:FR for across early timepoints.

Figure S11: The comparison of hormone induced gene expression changes against FR without IAA or GA-related genes.

## Acknowledgements

We thank Utrecht Sequencing Facility (Useq) providing illumina sequencing service for this research and Amber van Seters for her help with the IAA extractions.

## Author contributions

KK, RP and CKP: Conceptualization and Supervision.

LL: Funding Acquisition, Data Curation, Formal Analysis, Visualization, Writing – Original Draft.

LL, WK, TH, JW, GA, YK and CKP: Investigation.

LL, KK, RP, CKP, WK: Writing – Review & Editing.

All authors read and have approved the final manuscript.

## Data availability

The RNA sequencing data from this study are openly available in NCBI GEO repository reference number GSE255611.

